# Analysis of cortical dysplasias using b-tensor encoding diffusion MRI in an animal model

**DOI:** 10.1101/2025.11.19.689314

**Authors:** Olimpia Ortega-Fimbres, Ricardo Ríos-Carrillo, Edith Gaspar-Martínez, Priscila Ruiz-Acosta, Mirelta Regalado, Hiram Luna-Munguía, Alonso Ramírez-Manzanares, Luis Concha

**Affiliations:** Instituto de Neurobiología, Universidad Nacional Autónoma de México, Blvd. Juriquilla 3001, Querétaro, 76230, Querétaro, México.; Centre for Functional and Metabolic Mapping, Robarts Research Institute, Western University, 1151 Richmond Street, London, N6A 3K7, Ontario, Canada.; Departamento de Ciencias de Computación, Centro de Investigación en Matemáticas, A.C., Jalisco S/N, Guanajuato, 36023, Guanajuato, México.

**Keywords:** diffusion MRI, b-tensor encoding, cortical dysplasia, cortical microstructure

## Abstract

Cortical dysplasias are malformations of cortical development characterized by disorganization of the cyto- and myeloarchitecture of the neocortex. They are a common cause of epilepsy and their diagnosis through conventional imaging can often be challenging, hindering surgical treatments. Diffusion-weighted magnetic resonance imaging (dMRI) has the ability to infer tissue properties at the microscopic scale, making it a promising technique for detection of cortical dysplasias. This study aims to assess the microarchitecture of the cerebral cortex in a murine model of cortical dysplasia using diffusion-weighted magnetic resonance imaging (dMRI) acquired with b-tensor encoding. Pregnant Sprague-Dawley rats were administered either carmustine (BCNU) or saline solution on day 15 of gestation. Their offspring were imaged at 120 days of age using a 7 tesla scanner, acquiring diffusion-sensitive images with b-tensor encoding. Images were processed with Q-space trajectory imaging with positivity constraints (QTI+) to derive various metrics along a curvilinear coordinate system across the neocortex. After scanning, the brains were processed for immunofluorescence and histological examinations. Experimental animals exhibited a significant reduction of microscopic fractional anisotropy (µFA) and anisotropic kurtosis (Ka) in the middle and lateral cortical layers compared to the control animals. Immunofluorescence and histological analysis showed decreased and dysorganized myelinated fibers, and an increase of glial processes in BCNU-treated animals. Given the applicability of b-tensor encoding in clinical scanners, this approach holds promise for improving detection of focal cortical dysplasias in patients with epilepsy.

## Introduction

The cerebral cortex exhibits a highly organized architecture. Alterations during prenatal neurodevelopment can lead to diverse anatomical abnormalities that vary in extent and severity as a consequence of the precipitating insult. Focal cortical dysplasias (FCDs) are a specific type of malformation of cortical development, representing the first and second cause of pharmacoresistant focal epilepsy in children and adults, respectively (Blumcke et al. 2017). These malformations are characterized by disrupted cortical layers, neuronal heterotopia, and presence of dysmorphic neurons (Guerrini and Barba 2021). Magnetic resonance imaging (MRI) is the primary diagnostic tool used for detection of FCDs and to guide surgical interventions, when appropriate (Bernasconi et al. 2011). Diagnosis of FCDs can be challenging, as their detection relies on subtle visual cues such as focal cortical thickening, slight hyperintensity, and blurring of the gray/white matter boundary on conventional T1- or T2-weighted MRI (Walger et al. 2025). Moreover, there can be ample variation in their size and anatomical location (Colombo et al. 2009; Blackmon et al. 2014; Lee 2016). This results in frequent underdiagnosis and inadequate treatment, highlighting the need for more sensitive and specific imaging methods (Walger et al. 2025). Detection of FCDs can be improved substantially through quantitative methods that augment the diagnostic yield of conventional MRI, such as texture analysis (Bernasconi et al. 2001), voxel-based morphometry (Colliot et al. 2006), and artificial intelligence (Gill et al. 2021; Spitzer et al. 2022), and more recently through acquisition and analysis of MRI fingerprinting (Su et al. 2024; Ding et al. 2025).

Given that anatomical abnormalities that usually accompany FCDs can be subtle, there is a need for imaging methods that are able to capture the histopathological features that characterize these lesions. Diffusion-weighted MRI (dMRI) offers an alternative, non-invasive approach for studying tissue microarchitecture by measuring the diffusion of water molecules in different tissues (Beaulieu 2002; Concha 2014; Ghaderi et al. 2025). This technique can capture structural details that are not visible with conventional MRI, making it potentially more effective for diagnosing FCDs. Furthermore, dMRI can be applied to both human and animal models, enabling the translation of experimental findings to clinical practice (Alexander et al. 2017; Zhu et al. 2025). Several methods to analyze the diffusion signal have been introduced, the majority of which are implemented on data acquired using single diffusion encoding (SDE) (Stejskal and Tanner 1965). While these methods have been mainly used to characterize white matter, SDE dMRI can also provide relevant information regarding the laminar and columnar structure of the human, non-human primate, and rodent neocortex (Aggarwal et al. 2015; Feizollah and Tardif 2025; Leuze et al. 2014; McNab et al. 2013; Reveley et al. 2024, 2022; Villaseñor et al. 2023). Recent advances in dMRI acquisition and analysis methods have further improved the diagnostic capacity of this technique by providing more information on tissue characteristics (Topgaard 2019; Westin et al. 2016). Multidimensional encoding techniques have been developed to allow a more thorough analysis of brain microstructure by encoding diffusion through complex gradient waveforms (Lundell et al. 2019; Szczepankiewicz et al. 2021). In this sense, the b-tensor generalizes the scalar b-value and describes the shape, size and orientation of diffusion encoding, enabling interrogations of linear, planar, or spherical diffusion within the tissue. While diffusion tensor imaging (DTI) can only provide a single (average) diffusion tensor per voxel, and SDE results in biased estimates of microscopic anisotropy (Henriques et al. 2019), b-tensor encoding allows the construction of intra-voxel diffusion tensor distributions (DTD); this will provide rich information that better accounts for the heterogeneity of nervous tissue components (Magdoom et al. 2021). These innovations make b-tensor encoded dMRI a promising tool for characterizing the microarchitecture of the cortex and alterations present in FCDs and other cortical malformations (Nasser et al. 2022). In this study, we used advanced dMRI techniques to characterize tissue properties of the cortex in an animal model of FCD. Histological analyses were performed to bridge tissue properties with water diffusion patterns in the regions of interest. The aim was to assess whether b-tensor encoded dMRI methods are sensitive to the histopathological features of FCD as an initial exploration of their clinical applicability.

## Methods

All experimental procedures were approved by the Ethics Committee of the Institute of Neurobiology (protocol 111-A).

### Murine Model of Cortical Dysplasia

To induce the histopathological features of FCD Type IIa in rodents, we used a known animal model that disrupts corticogenesis *in utero*. Pregnant Sprague-Dawley rats were injected with a single dose of either the alkylating agent BCNU (bis-chloroethylnitrosourea, also known as carmustine; 20 mg/kg in saline solution, i.p.), or saline solution (for control) on embryonic day 15; a time point that corresponds to the peak of cortical neurogenesis (Benardete and Kriegstein 2002). Based on previous reports (Inverardi et al. 2013; Moroni et al. 2008) and our observations (Aquiles et al. 2023; Villaseñor et al. 2023), this procedure is known to induce cortical alterations in the pups similar to those found clinically (Blumcke et al. 2017; Najm et al. 2022). The pups were kept with their mothers until weaning. All animals had access to food and water *ad libitum* and were always kept at the animal facility under controlled environmental conditions. Experiments were only conducted with the offspring. A total of 18 control (6 female) and 20 BCNU-treated (8 female) rats were included for further analysis.

### Diffusion-Weighted Magnetic Resonance Imaging

Images were acquired at the National Laboratory for Magnetic Resonance Imaging (Lanirem) in Juriquilla, Queretaro, Mexico, using a 7 T Bruker Pharmascan pre-clinical MRI scanner and a 2×2 array head surface coil. The rats were anesthetized with isoflurane (4% for induction, 2% for maintenance) and kept warm by circulating water at 37 °C through hoses placed under the scanner bed. Vital signs were continuously monitored throughout the study using a compatible system. A single imaging session was performed for each animal (four months old) to determine diffusion parameters in the dysplastic and normal cortices. Each session lasted one hour. dMRI were obtained using an open-source sequence based on a 2D spin-echo echo-planar acquisition sequence, available from the Preclinical Neuro MRI repository (https://github.com/mdbudde/mcw_Preclinical_MRIsequences). Images with coronal orientation were acquired with voxel resolution of 200×200×1010 µm^3^, repetition time (TR) = 2000 ms, echo time (TE) = 40.86 ms, flip angle = 90°. The implemented protocol consisted of three b-tensor shapes: linear, spherical and planar (Szczepankiewicz et al. 2019). The spherical diffusion encoding gradients (SDE) were optimized prior to acquisition by using the NOW toolbox (https://github.com/jsjol/NOW) (Sjölund et al. 2015) and scaled in magnitude to obtain four b-values (200, 700, 1400, and 2000 s/mm^2^). To retain gradient spectral characteristics between waveforms, the planar and linear tensor encoding gradients (LTE and PTE, respectively) were extracted from the optimized SDE, using one axis for LTE and the other two for PTE (Lundell et al. 2019). The STE waveform was rotated in 10 directions at each b-value; LTE and PTE waveforms were rotated to obtain [10, 10, 16, 46] directions for each corresponding b-value. Supplementary Figure S1 shows the waveforms and protocol used in this study.

### DWI processing

The images were preprocessed to minimize noise and artifacts. This included noise reduction (Cordero-Grande et al. 2019) and correction of geometric inhomogeneities (Andersson and Sotiropoulos 2016) using Mrtrix 3.0.4 (Tournier et al. 2019) and fsl 6.0.7.1 (Smith et al. 2004) (Fig. 1A). Q-space trajectory imaging with positivity constraints (QTI+) (Herberthson et al. 2021) was computed as implemented in https://github.com/DenebBoito/qtiplus. In addition to the four diffusion-tensor metrics, namely fractional anisotropy (FA), axial, radial and mean diffusivities (AD, RD and MD, respectively) (Fig. 1B, top panel), QTI+ provides four novel metrics: microscopic anisotropy (µFA), microscopic orientation coherence (CC), and anisotropic and isotropic kurtosis (K_a_, and K_i_, respectively) (Westin et al. 2016) (Fig. 1B, bottom panel). These summary metrics simplify the interpretation of the distribution of tensors of different shapes, sizes, and orientations, and are sensitive to features of microarchitecture (Topgaard 2019). The valid range for FA, µFA and CC was between 0 and 1, and voxels outside of these ranges (less than 0.5% of all voxels) were attributed to fitting errors and excluded from further analyses.

**Fig. 1.**
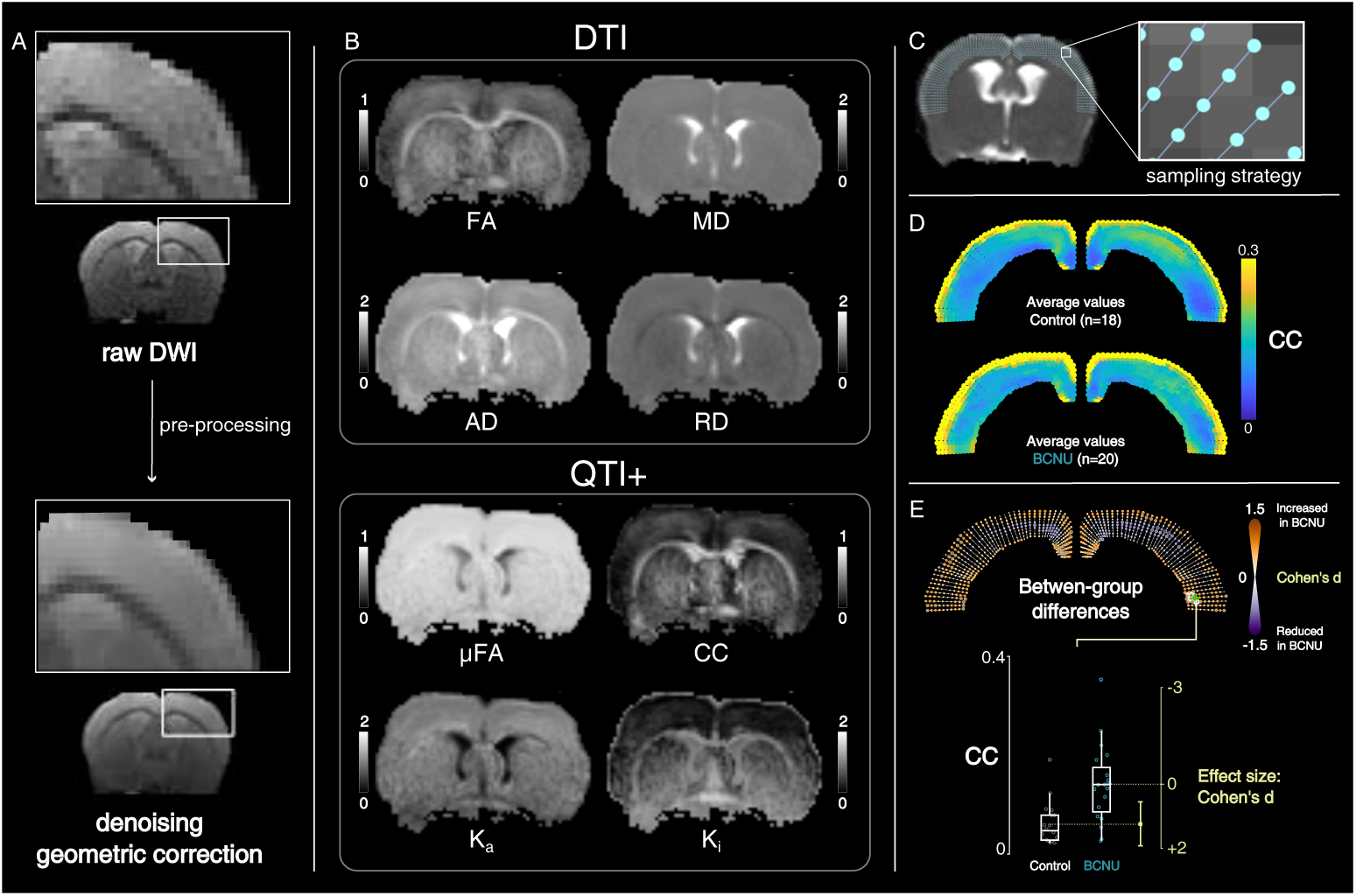
Analysis pipeline. A: Raw dMRI were preprocessed to minimize noise and correct for geometric inhomogeneities. B: QTI+ provided four DTI metrics (top): fractional anisotropy (FA), and axial, radial and mean diffusivities (AD, RD, MD, respectively; in units of mm^2^*/*s), as well as (bottom) microscopic anisotropy (µFA), microscopic orientation coherence (CC), and anisotropic and isotropic kurtosis (K_a_ and K_i_, respectively). C: A curvilinear coordinate system (50 gridlines spanning medial to lateral, and 10 depth levels from the pial border to the gray/white matter border) was defined in each rat brain, providing inter-subject anatomical correspondence. Descriptive statistics were obtained from the data sampled by the gridlines, the point-wise average values are shown for each group (D). Student’s *t*-tests were subsequently performed at each point followed by correction for multiple comparisons (E). Marker size and color indicate effect size (Cohen’s *d*), gray circles indicate point-wise p_uncorr_<0.01, white areas indicate significant clusters (p_clus_<0.05). A Gardner-Altman plot presents data from one exemplary vertex (green circle in Cohen’s *d* map).

### Spatial Analysis

A curvilinear coordinate system with anatomical correspondence between animals was constructed for each rat to sample dMRI parameters across the depth and extent of the cortex (Villaseñor et al. 2023). For this purpose, in a single coronal slice (located approximately -0.8 mm posterior to bregma, harboring primary motor and somatosensory cortices [M1 and S1, respectively]), the pial surface of the brain and the boundary between gray and white matter were manually delineated. A Laplacian potential field was simulated between these two boundaries (Lerch et al. 2008). From the pial surface, 50 evenly distributed virtual trajectories (gridlines) were created and propagated organically through the Laplacian field; each trajectory extended towards the white matter boundary, following the curvature of the cortex in a manner analogous to cortical columns. The dMRI parameters were sampled at 10 equidistant points along each of these gridlines (Fig. 1C). The code to create this curvilinear grid is available at https://github.com/lconcha/Displasias.

### Statistical Analysis

Acknowledging that cortical cyto-and myeloarchitecture varies among cortical regions (Kleven et al. 2023), the analysis was conducted in a spatially dependent fashion. Abnormalities were assessed separately for each dMRI metric using univariate statistics (Fig. 1D,E). At each point in each gridline, Student’s *t*-tests were performed to detect differences between the two groups (Fig. 1E). Effect size estimation (the magnitude of the difference between groups) was calculated using Cohen’s *d*. Statistical analyses were corrected to minimize the probability of Type I errors (false positives) by using cluster-level permutation tests (Cox et al. 2017). The vertex-wise cluster-forming threshold was set as p_uncorr_<0.01. Cluster significance (p_clus_) was determined by randomizing the data between experimental groups, performing 5,000 permutations to obtain the distribution of cluster sizes that could arise by chance. From this empirically-derived null distribution, the probability of finding clusters with a similar extent to those observed in the real data (i.e., without randomization between groups) was calculated. Cluster-wise statistical significance was defined as p_clus_<0.05

## Histological Analysis

After completing the dMRI studies, all animals were intracardially perfused with 0.9% NaCl solution followed by 4% paraformaldehyde (PFA) solution. The brains were removed and preserved in fresh PFA 4% solution for 24 h at 4 °C. After this, each brain was immersed in a 20% sucrose solution for 48 h, followed by a 30% sucrose solution for another 48 h. Brains were stored at -72 °C until further analysis. Coronal sections (20 µm-thick) from the region of interest were obtained using a cryosotat (Leica) based on the following Paxinos and Watson Rat Brain Atlas interaural coordinates: 8.74-8.08 mm. Slices were kept in a cold 1X phosphate buffer solution (PBS; Sigma-Aldrich). Immunofluorescence was performed using the primary antibodies anti-Myelin Basic Protein (MBP; 1:500; abcam), anti-Neuronal Nuclear Protein (NeuN; 1:350; abcam), and anti-Glial Fibrillary Acidic Protein (GFAP; 1:350; Sigma-Aldrich). For the triple immunofluorescence staining, the tissue sections were blocked with Bovine Serum Albumin (BSA; Sigma-Aldrich) 2% solution + 0.3% triton X-100 (ThermoFisher) in 1X PBS for 45 min. The sections were incubated with the primary antibodies (MBP and NeuN) for 24 h at 4°C. Then, slices were washed five times for 10 min in PBS 1X + Tween 0.1% solution (Sigma-Aldrich). Secondary antibodies conjugated with fluorescent dyes (AlexaFluor, goat anti-mouse-647 and goat anti-rabbit-555) were diluted 1:500 in a solution of PBS 1X + 0.1% Tween and incubated for 4 h at 4°C. After incubation, the slices underwent five washes (each one lasting 10 min) in PBS 1X. The sections were then blocked with a solution of BSA 2% + 0.3% Triton X-100 in PBS 1X for 45 min. Finally, GFAP was added in a PBS 1X + 0.1% Tween solution and incubated for 24 h at 4°C. Afterwards, another set of five 10-min washes in PBS 1X + 0.1% Tween was done, and the corresponding secondary antibody was added (AlexaFluor, goat anti-mouse-488) for 4 h, followed by five additional washes in 1X PBS. Finally, the slices were mounted using Mowiol. Brain slices were imaged using a confocal microscope (Zeiss LSM 880, with 488/594/647 nm wavelengths) and a fluorescence microscope (Zeiss Apotome, with 488/594 nm wavelengths). The system of this last microscope was linked to a computer running AxioVision software (version 4.8), where the MosaiX module was used to acquire mosaic images at 10X.

The photomicrographs were evaluated using Fiji (Schindelin et al. 2012). Samples were taken from the primary motor cortex (M1) and the primary somatosensory cortex (S1). In these samples, the brightness threshold was automatically adjusted (Li and Tam 1998), and the spatial profile of glial density was also determined by calculating the percentage of the area occupied by the cells. The organization of the myeloarchitecture was evaluated through structure tensor analysis (Budde and Frank 2012), as implemented in OrientationJ (https://github.com/Biomedical-Imaging-Group/OrientationJ) (Püspöki et al. 2016), calculating vector and local coherence maps using a Gaussian window of 15 µm.

## Results

### Analysis of Diffusion-Weighted Magnetic Resonance Images

All diffusion metrics showed spatial variability across the extent and depth of the cortex, which is not captured by histogram analyses of the entire cortex (Fig. 2 and 3). In control animals, the middle layers of the cortex showed the highest FA and lowest RD in the middle layers of M1 and S1 regions. Contrarily, the most lateral aspects of the cortex showed the lowest average FA values. Mean and axial diffusivities were less heterogeneous across the cortex, with the exception of the most superficial layers. Metrics derived from QTI+ in control animals showed very high values of µFA in the middle layers of the cortex with decreasing values towards the lateral aspects of the cortex. This pattern was mirrored by K_a_. A band of very high CC and K_i_ values was seen in the most superficial layers of the cortex. Average between-group differences are readily visible for µFA, K_a_, and K_i_ in the spatial maps and in the histogram analyses. BCNU-treated animals showed decreased FA values in the central-medial areas of the cortex in both hemispheres, encompassing the motor cortices M1 and M2, the primary somatosensory, and the cingulate cortex (Fig. 2). This reduction of FA was accompanied by an increase of axial diffusivity and a reduction of axial diffusivity that was significant at the cluster level in the left hemisphere. Mean and axial diffusivities did not show any statistically significant differences between groups. QTI+ metrics attainable only through b-tensor encoding provided additional information (Fig. 3). Both microscopic anisotropy and anisotropic kurtosis showed extended reductions along the cortex of both hemispheres in BCNU-treated animals (including cingulate, motor, and somatosensory cortex), mostly in the middle to superficial layers. The deep layers of the most lateral aspect of the left hemisphere showed an increased of CC, while the middle layers of the rest of the cortex showed reductions that were not significant at the cluster level. Finally, there were distributed increases of K_i_ throughout the superficial layers of the cortex in both hemispheres.

**Fig. 2.**
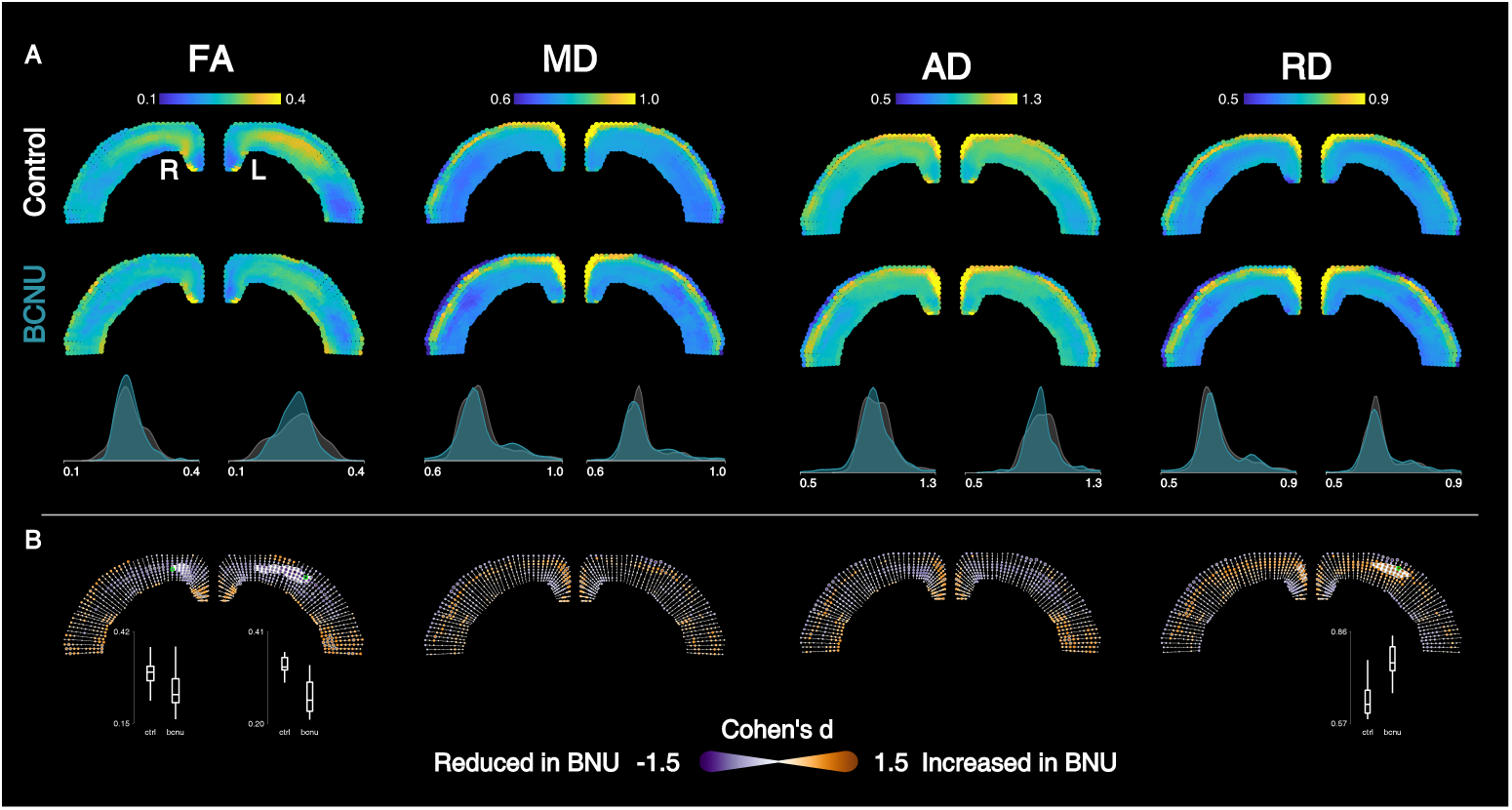
DTI metrics. A: Average values for each metric are shown for control and BCNU-treated animals (first two rows). Histograms show the group-wise average values across the entire cortex per hemisphere (third row). B: Between-group differences illustrated as in Fig. 1E. Effect sizes (Cohen’s *d*) are color-coded at each point per grid line (marker size represents |*d*|). Gray circles indicate vertex-wise p_uncorr_<0.01, and white areas indicate cluster-corrected statistical significance (p_clus_<0.05). Box plots of vertices identified in green are shown below for hemispheres with significant clusters.

**Fig. 3.**
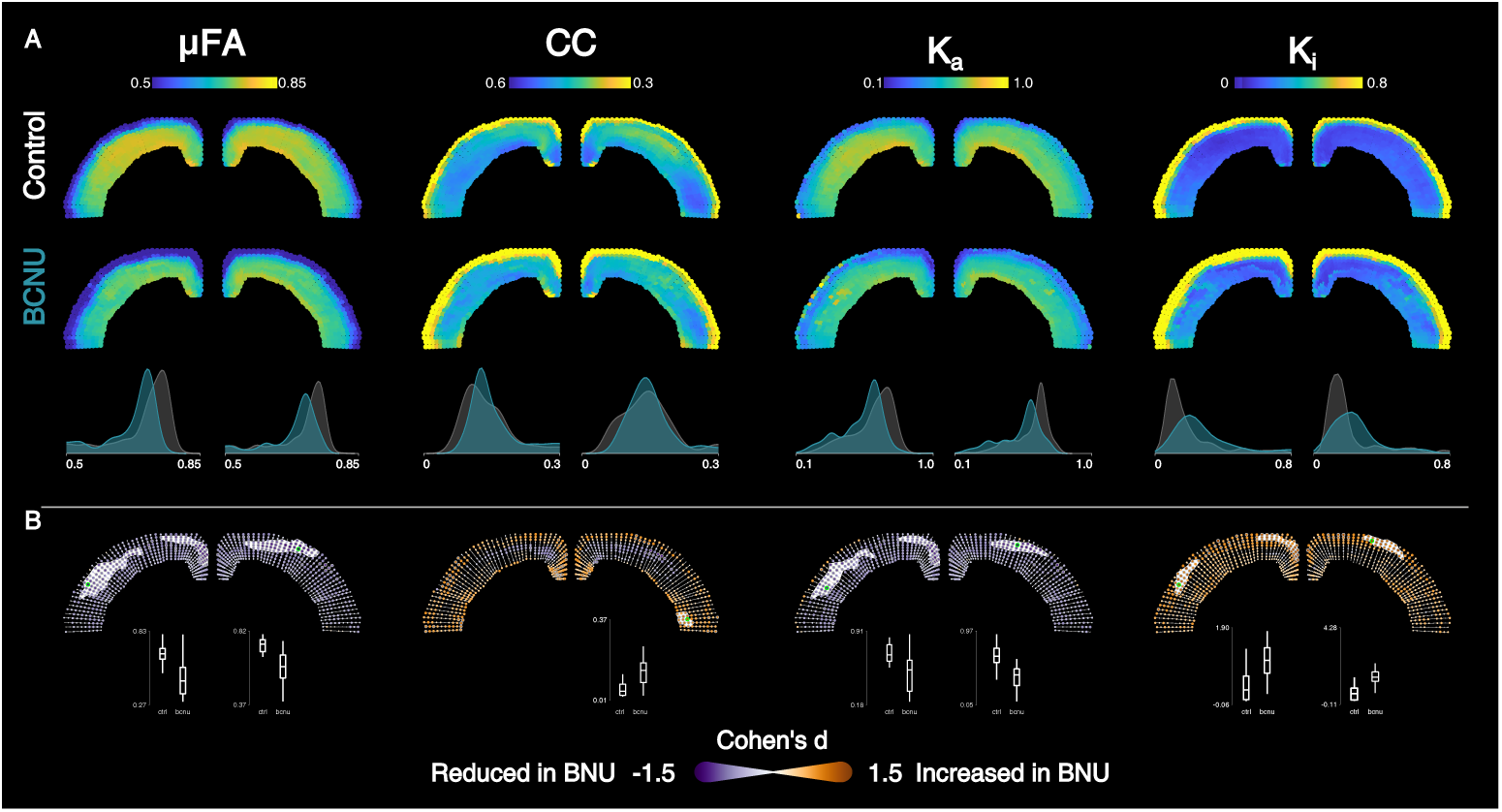
QTI+ metrics. Figure layout and description as in Fig. 2.

### Histology

Qualitative evaluations revealed reduced intracortical myelination in BCNU-treated rats, with reduced MBP+ fibers in the most superficial cortical layers. Additionally, variations in the distribution of neuronal nuclei (NeuN+) and morphological/quantitative changes in astrocytes were observed in BCNU-treated rats compared to the control group (Fig. 4).

**Fig. 4.**
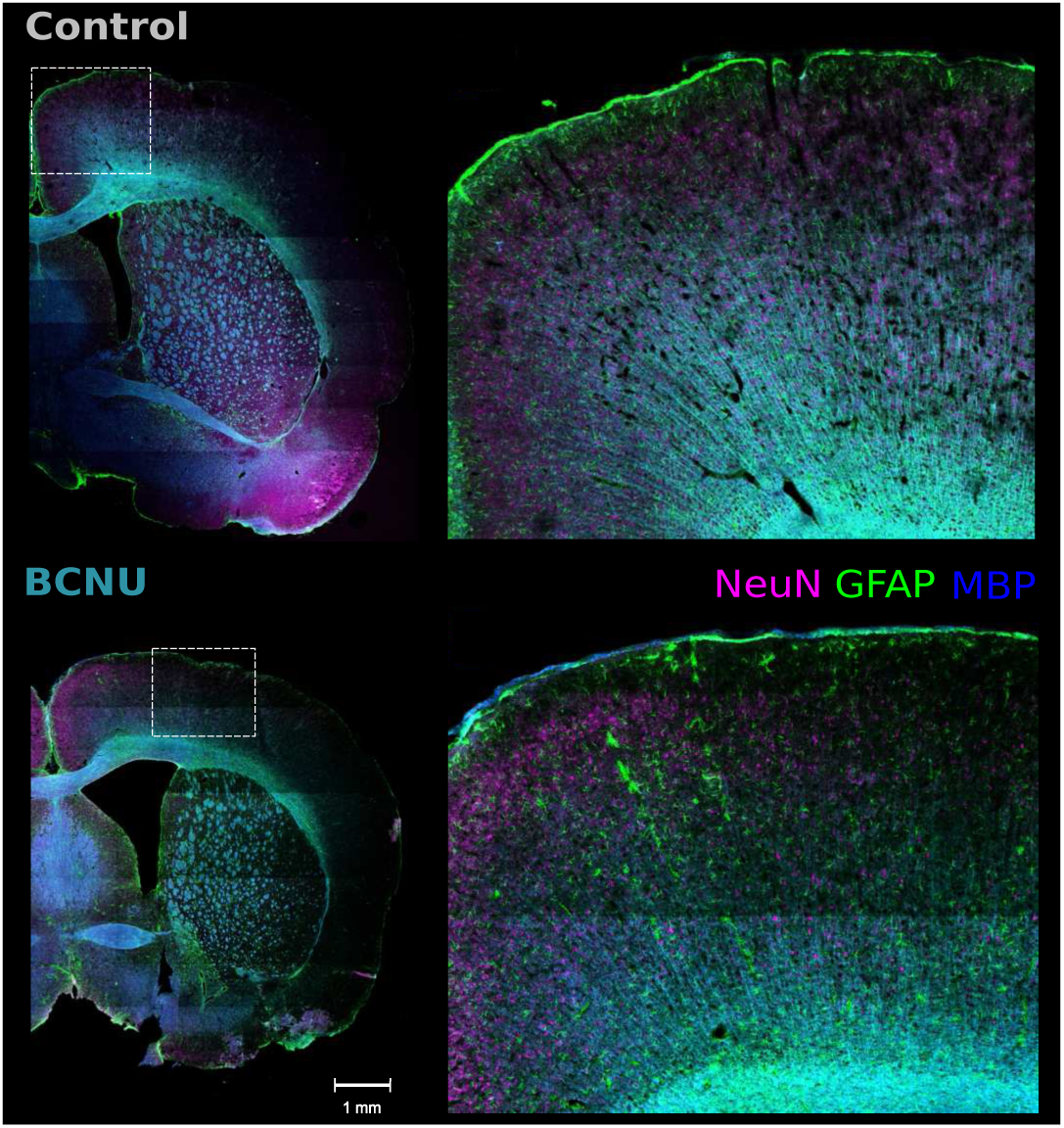
Triple-labelled Immunofluorescence showing neuronal nuclei (NeuN; in magenta, 594 nm), glia (GFAP; in green, 488 nm), and myelinated fibers (MBP; in blue, 647 nm) from a control (top) and a BCNU-treated rat (bottom). The experimental animal shows overall less intracortical myelin, heterogeneous spatial distribution of neuronal nuclei, and increased astrocytic processes.

### Myelin Analysis

Quantitative examination of histology through structure tensor analysis of MBP-labelled immunofluorescent photomicrographs revealed differences of the myeloarchitecture of the cerebral cortex between the control and experimental animals. BCNU-treated animals showed decreased coherence from the medial to the most lateral region of the cortex. This was further corroborated by vector maps, where interspersed vector vortices were observed in the somatosensory region, indicating a disorganization of myelin fibers (Fig. 5).

**Fig. 5.**
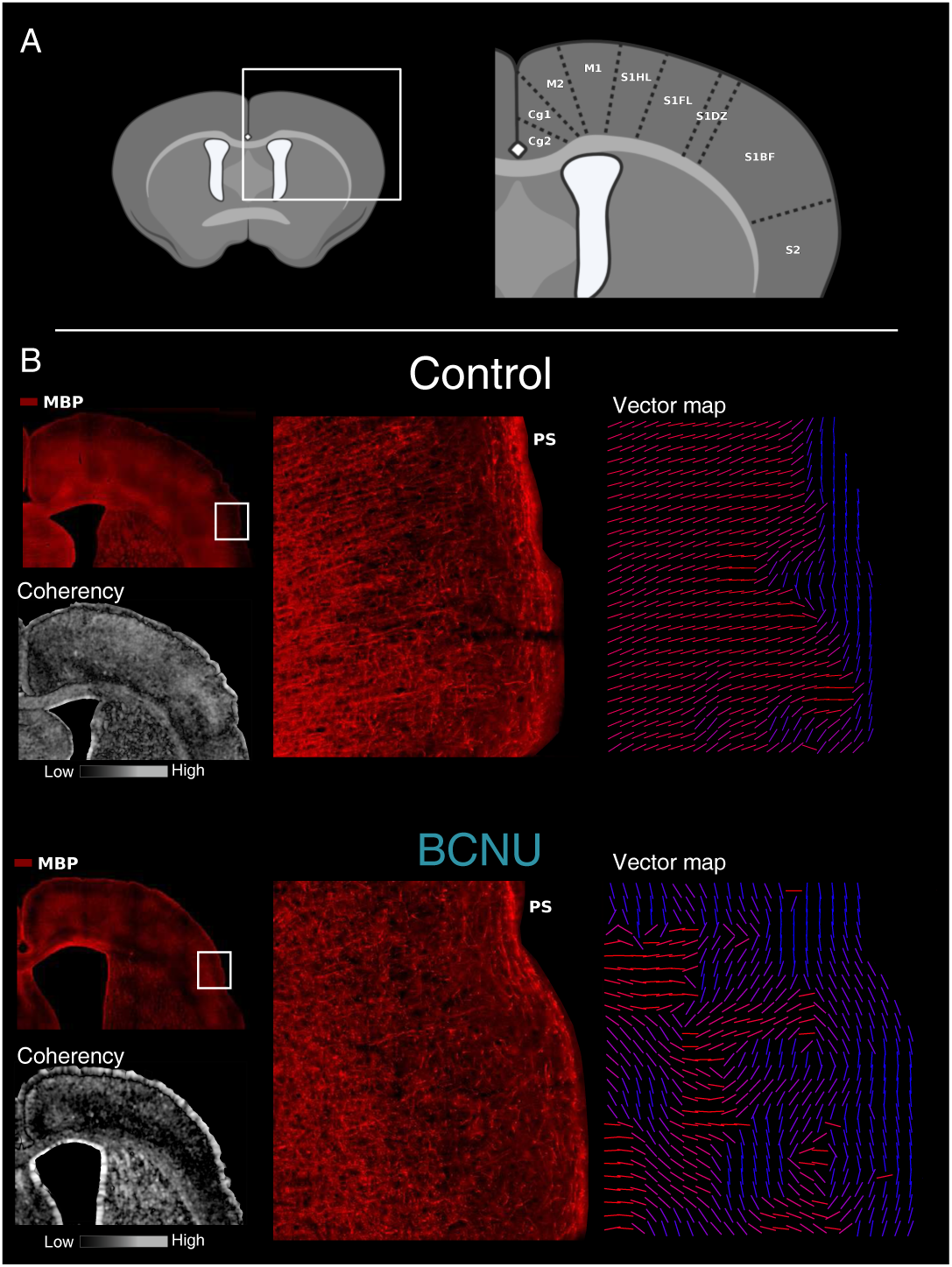
Structural Tensor Analysis of MBP+ fibers. A. Cortical regions covered during the analysis. B. Exemplary control (top) and BCNU-treated animals showing MBP+ fibers (red) and corresponding coherence maps (grayscale) derived from structure tensor analysis. Experimental animals show reduced coherence throughout the cortex, from the medial area (M2) to the somatosensory area (S2). On the right, enlarged sections encompassing S1BF with its corresponding vector maps indicating the local main orientation of MBP+ fibers (red: medial-lateral; blue dorsal-ventral orientations), where coherent directions are observed in the control group, and interspersed vortices in the experimental group. Cg1 and Cg2: cingulate cortices; M1 and M2: primary and secondary motor areas; S1, S1FL, S1DZ, and S1BF: primary somatosensory cortices (hindlimb, forelimb, dysgranular zone, barrel field); S2: secondary somatosensory cortex.

### Histological Analysis of Astrocyte Cortical Distribution

There was an overall increase of the percentage of area occupied by astrocytes (GFAP+) between the control and experimental groups in the primary motor cortex (Fig. 6, top row). Depth-wise analysis revealed increased GFAP+ area particularly in cortical layers IV-VI (Fig. 6, middle panel). This increased presence of glial processes was not as marked in the S1 region (Fig. 6, bottom row).

**Fig. 6.**
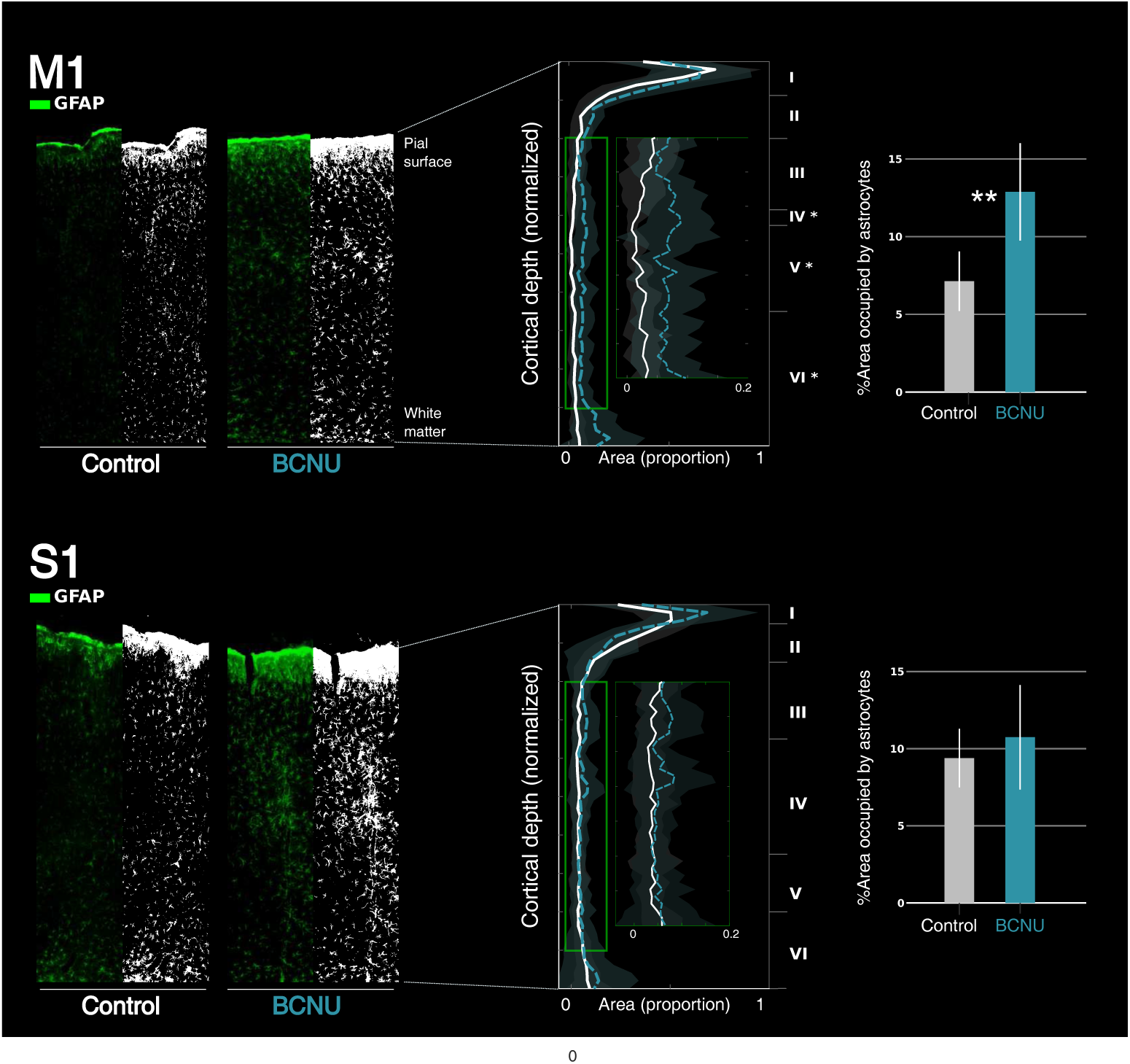
Quantification of area occupied by astrocytes (GFAP+) in the cortical regions M1 (top row) and S1 (bottom row). For each cortical region, leftmost panels show the GFAP immunofluorescence (green) in the control and BCNU groups, and the corresponding binarized image (black and white). Middle panels show the groupwise, depth-dependent spatial density profiles, with layers II-V enlarged in the insets. GFAP+ area is larger in experimental animals, particularly in layers IV-VI. Bar graphs in the rightmost panels show between group differences throughout the entire cortical region (**: p <0.01).

## Discussion

Despite the high spatial resolution attainable with modern anatomical MRI, identification of FCD in patients with focal-onset epilepsy remains a challenge, as their microscopic disarray can hide behind tissue that looks deceptively ordinary at macroscopic scales. In this work we show that advanced dMRI with b-tensor encoding is able to extract information at the mesoscopic level, evidencing subtle histopathological landmarks characteristic of cortical malformations.

We used an animal model that presents the subtle histological cyto- and myeloarchitectonic irregularities characteristic of human FCD (Aquiles et al. 2023; Benardete and Kriegstein 2002; Villaseñor et al. 2023). In prior work from our group using the same model we showed that multi-tensor fit of the diffusion signal was able to separate groups of intracortical fibers depending on their orientation to the cortical surface (i.e., radial and tangential fibers) and, moreover, differentially identify diffusion abnormalities in BCNU-treated animals (Villaseñor et al. 2023). Here, we used QTI+ to derive diffusion tensor metrics, and found reductions of FA in the middle cortical layers, in close agreement with our previous report, despite the age difference of the rats between the two studies (P30 vs P120). These results are in line with previous reports of reduced FA within FCD lesions (Gennari et al. 2023; Gross et al. 2005) and also in the superficial white matter adjacent to the lesions (Lee et al. 2004; Lorio et al. 2020; Urquia-Osorio et al. 2022). DTI is a commonplace method, available in virtually all MRI scanners. The robustness of the FA findings across studies indicates that this metric has the potential to aid in the identification of FCD in patients (Madan and Grant 2009). Its sensitivity, however, is likely reduced as a consequence of the known limitations of DTI that render it insufficient to characterize the complex organization of the neocortex. Previous reports have shown the utility of diffusion metrics derived from neurite orientation dispersion and density imaging (NODDI) (Zhang et al. 2012), such as microscopic anisotropy and intracellular and intraneurite volume fractions, for the detection of FCD (Lorio et al. 2020; Rostampour et al. 2018; Winston et al. 2014). Notably, NODDI is specifically designed to characterize white matter, and therefore may be unfit for the study of the neocortex. Extending the ideas of NODDI, soma and neurite density imaging (SANDI) provides an approximation to the biophysical properties of gray matter with the inclusion of a sphere compartment to model cell bodies (Palombo et al. 2020). Metrics derived from SANDI are good indicators of gray matter microstructure in animal models (Ianuş et al. 2022), as well as in humans (Barakovic et al. 2024; Genc et al. 2025; Lee et al. 2024), and therefore constitute a viable option for the detection of FCD in future studies.

The SDE method (Stejskal and Tanner 1965) is extremely useful for the acquisition of DWI and their subsequent analyses through many different methods. Nonetheless, novel acquisition schemes with free gradient waveforms sample the signal more richly across measurement space (Westin et al. 2016). From these, novel and complementary diffusion metrics tailored to separate microscopic anisotropy from orientation dispersion and other sources of variance can be obtained through the examination of diffusion tensor distributions (Herberthson et al. 2021; Topgaard 2019; Westin et al. 2016). Acquiring independent observations in a multidimensional manner enhances the characterization of heterogeneous media (Lundell et al. 2019). In our work, QTI+ metrics showed reductions of microscopic anisotropy that were more extensive than the corresponding DTI abnormalities, as evidenced by the two correlated metrics µFA and K_a_ (the former being the normalized value of the latter). These reductions can be interpreted as compromised axonal structure (Hansen 2019; Li et al. 2021; Shemesh 2018). In parallel, K_i_ showed an overall increase in the superficial layers of the cortex in BCNU animals, concordant with gliosis and heterogeneity of cellular size and morphology (Glenn et al. 2015; Hui et al. 2015). Our histological examinations, as well as prior evaluations of glial cells in BCNU-treated animals (Rodríguez-Arzate et al. 2021) support this interpretation. The utility of b-tensor encoding for the detection of different forms of malformations of cortical development has been explored (Lampinen et al. 2020), with findings in FCD that are in line with our results. The added value of b-tensor encoding and QTI+, together with the feasibility to perform this type of acquisition efficiently in the clinic (Nilsson et al. 2020) open new avenues for the detection of FCD.

The animal model used here shows histopathological features similar to those observed clinically (Najm et al. 2022), including cortical dyslamination and disarray of the myeloarchitecture (Aquiles et al. 2023; Inverardi et al. 2013; Villaseñor et al. 2023). Reductions in microscopic orientation coherence (CC) were observed in the middle cortex of BCNU-treated animals (Fig. 3) and are likely associated with the altered geometric disposition of myelinated fibers (Fig. 4 and 5), despite not reaching cluster-level significance. Of note, an increase in glial processes was observed in BCNU-treated animals compared to controls, especially in layers IV-VI of M1 region (Fig. 4 and Fig. 6). Contrastingly, S1 showed no significant astrocytic density change. Differences of cortical architecture may render M1 more susceptible to microstructural disruption (Benedetti et al. 2020; Lee et al. 2022). Prominent gliosis due to oxidative stress and inflammation is a common finding in human dysplastic cortex and in many non-genetic animal models (Arena et al. 2019; Kakita et al. 2005; Kielbinski et al. 2016). While ongoing seizures are a cause of reactive gliosis, astrocytic gliosis alone is capable of initiating epileptic activity (Robel et al. 2015). Identification of focal gliosis is, therefore, another opportunity for the detection of FCD, and has in fact been explored with, for example, radiotracers (Butler et al. 2013) and manganese-enhanced MRI (Kawai et al. 2010). In agreement with other reports that used dMRI to detect gliosis (Benjamini et al. 2023; Chary et al. 2023; Garcia-Hernandez et al. 2022), our QTI+ results are indicative of this possibility.

Our study has limitations to consider. While the animal model used here induces alterations of cortical microarchitecture typical of FCD, it does not encompass the range of abnormalities seen in humans. Notably, the cortex of BCNU-treated animals lacks balloon cells, a prominent feature of FCD Type IIb; consequently, the model used more closely resembles FCD Type IIa (Blümcke et al. 2021). Therefore, we cannot ascertain the impact that such abnormal cells may have on the diffusion signal, nor if milder alterations of cyto- and myeloarchitecture characteristic of FCD Type I are enough to alter diffusion metrics. Evaluation of the sensitivity and specificity of diffusion metrics to detect FCD would require a gold standard to be compared to. However, the cortical abnormalities induced by BCNU are not focal but rather distributed across the cortex, which contrasts to the moderately well-demarcated FCD lesions observed in humans. Other animal models, such as freeze lesions of the cortex, induce focal alterations (Lau and Dulla 2017), but can include macroscopic abnormalities and the creation of a microgyrus (Wong 2009). Such gross anatomical alterations are easily identified with conventional MRI, thus rendering dMRI unnecessary, and are against our ultimate goal to detect the subtle mesoscopic abnormalities that often go unnoticed in patients. While all animals were processed for histology, technical difficulties during tissue handling preclude a one-to-one comparison of dMRI and microphotographs, and the evaluation of correlations between diffusion metrics and quantitative histology. Diffusion time dependence and water exchange effects, which may modulate QTI+ and other dMRI metrics (Novikov et al. 2016), were not accounted for in this study. This effect may be even more relevant in gray matter given its abundance of non-myelinated (and therefore more water-permeable) axons and dendrites (Dong et al. 2025; Lee et al. 2020; Olesen et al. 2022). Future studies would benefit from this addition to b-tensor encoding. Finally, while b-tensor encoding is feasible in clinical scanners (Nilsson et al. 2020), spatial resolution will be an important hurdle to overcome in order to provide sufficient sampling of the human cortical mantle, which typically has a thickness of 1.5-4.5 mm (Fischl and Dale 2000), thus making super-resolution techniques highly desirable (Vis et al. 2021).

Our imaging and histology findings paint a coherent picture that highlights the ability of b-tensor encoding to capture the subtle histopathological abnormalities present in FCD. QTI+ resolved fine-grained attributes (microstructural anisotropy loss, orientation dispersion, and intra-voxel heterogeneity) closely tracking myelin disorganization and glial changes. The benefits of b-tensor encoding and QTI+ are especially valuable in the cortex, where the complex architecture of the tissue poses challenges to simpler diffusion models.

## Acknowledgements

The authors thank Dr. Juan Ortiz-Retana for assistance during MRI acquisition; Drs. José Martín García Servín, Alejandra Castilla León and María A. Carbajo Mata, for their help at the animal facility. We thank the personnel at the histology facility, particularly Nydia Hernández-Ríos, María Lourdes Palma-Tirado, and Ericka Alejandra De los Ríos; Drs. Remy Fernand Avila Foucat and Reinher Pimentel-Domínguez provided further assistance for immunofluorescence. Image processing was partially performed using the National Laboratory for Advanced Scientific Visualization (LAVIS) with help from Luis Aguilar and Alejandro de León. Additional computing assistance was provided by Leopoldo González-Santos.

## Funding

Conahcyt/Secihti (CF-2023-I-218 LC) and UNAM-DGAPA (IN204720 and IN213423 LC, IN211326 HLM). The first author received a scholarship from SECIHTI (1019538).

## Supporting information

**Supplementary Figure S1.**
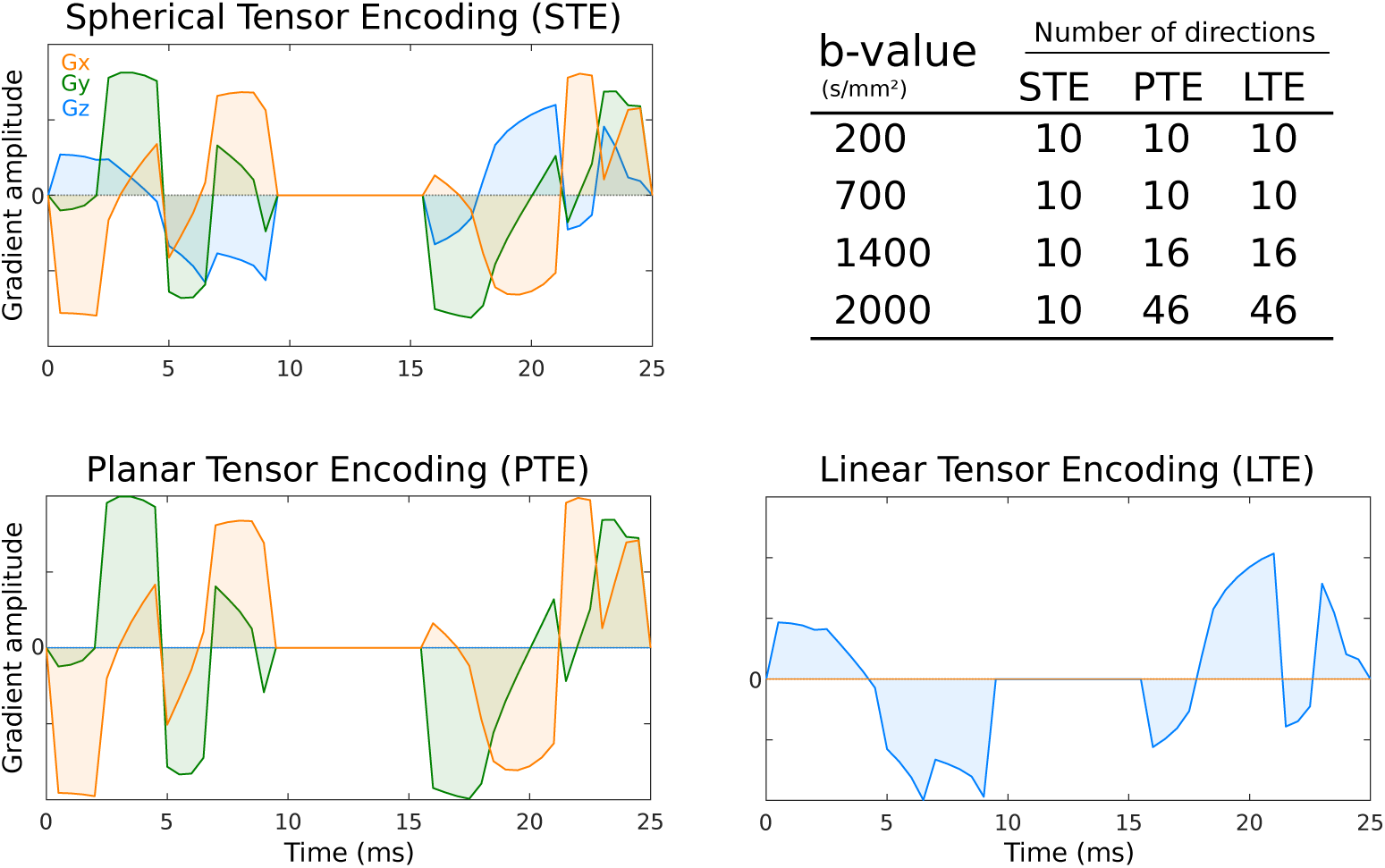
Diffusion gradient waveforms. Three exemplary waveforms are shown for spherical, planar, and linear tensor encodings (STE, PTE and LTE, respectively). STE waveforms were optimized then used as a base to create PTE waveforms from its Gx and Gy components, and LTE waveforms using Gz. Gradient amplitude (mT/m) was scaled to create different b value scalings. The table at the top right shows the number of directions for each b-tensor shape and b value scaling.).

## Notes

### Competing Interest Statement

The authors have declared no competing interest.

## References

Aggarwal M, Nauen DW, Troncoso JC, et al (2015) Probing region-specific microstructure of human cortical areas using high angular and spatial resolution diffusion mri. NeuroImage 105:198–207. doi:10.1016/j.neuroimage.2014.10.053

Alexander DC, Dyrby TB, Nilsson M, et al (2017) Imaging brain microstructure with diffusion mri: practicality and applications. NMR in biomedicine doi:10.1002/nbm.3841

Andersson JLR, Sotiropoulos SN (2016) An integrated approach to correction for off-resonance effects and subject movement in diffusion mr imaging. NeuroImage 125:1063–1078. doi:10.1016/j.neuroimage.2015.10.019

Aquiles A, Fiordelisio T, Luna-Munguia H, et al (2023) Altered functional connectivity and network excitability in a model of cortical dysplasia. Scientific Reports 13(11):12335. doi:10.1038/s41598-023-38717-2

Arena A, Zimmer TS, van Scheppingen J, et al (2019) Oxidative stress and inflammation in a spectrum of epileptogenic cortical malformations: molecular insights into their interdependence. Brain Pathology (Zurich, Switzerland) 29(3):351–365. doi:10.1111/bpa.12661

Barakovic M, Weigel M, Cagol A, et al (2024) A novel imaging marker of cortical “cellularity” in multiple sclerosis patients. Scientific Reports 14(1):9848. doi:10.1038/s41598-024-60497-6

Beaulieu C (2002) The basis of anisotropic water diffusion in the nervous system - a technical review. NMR in Biomedicine 15(7-8):435–55. doi:10.1002/nbm.782

Benardete EA, Kriegstein AR (2002) Increased excitability and decreased sensitivity to gaba in an animal model of dysplastic cortex. Epilepsia 43(9):970–982. 10.1046/j.1528-1157.2002.40901.x

Benedetti B, Dannehl D, Janssen JM, et al (2020) Structural and functional maturation of rat primary motor cortex layer v neurons. International Journal of Molecular Sciences 21(17):6101. doi:10.3390/ijms21176101

Benjamini D, Priemer DS, Perl DP, et al (2023) Mapping astrogliosis in the individual human brain using multidimensional mri. Brain 146(3):1212–1226. doi:10.1093/brain/awac298

Bernasconi A, Antel SB, Collins DL, et al (2001) Texture analysis and morphological processing of magnetic resonance imaging assist detection of focal cortical dysplasia in extra-temporal partial epilepsy. Ann Neurol 49(6):770–5. doi:10.1002/ana.1013

Bernasconi A, Bernasconi N, Bernhardt BC, et al (2011) Advances in mri for “cryptogenic” epilepsies. Nat Rev Neurol 7(2):99–108. doi:10.1038/nrneurol.2010.199

Blackmon K, Kuzniecky R, Barr WB, et al (2014) Cortical gray-white matter blurring and cognitive morbidity in focal cortical dysplasia. Cerebral Cortex p bhu080. doi:10.1093/cercor/bhu080

Blumcke I, Spreafico R, Haaker G, et al (2017) Histopathological findings in brain tissue obtained during epilepsy surgery. The New England Journal of Medicine 377(17):1648–1656. doi:10.1056/NEJMoa1703784

Blümcke I, Coras R, Busch RM, et al (2021) Toward a better definition of focal cortical dysplasia: An iterative histopathological and genetic agreement trial. Epilepsia 62(6):1416–1428. doi:10.1111/epi.16899

Budde MD, Frank JA (2012) Examining brain microstructure using structure tensor analysis of histological sections. NeuroImage 15(63):1–10. doi:10.1016/j.neuroimage.2012.06.042

Butler T, Ichise M, Teich AF, et al (2013) Imaging inflammation in a patient with epilepsy due to focal cortical dysplasia. Journal of Neuroimaging: Official Journal of the American Society of Neuroimaging 23(1):129–131. doi:10.1111/j.1552-6569.2010.00572.x

Chary K, Manninen E, Claessens J, et al (2023) Diffusion mri approaches for investigating microstructural complexity in a rat model of traumatic brain injury. Scientific Reports 13(1):2219. doi:10.1038/s41598-023-29010-3

Colliot O, Bernasconi N, Khalili N, et al (2006) Individual voxel-based analysis of gray matter in focal cortical dysplasia. Neuroimage 29(1):162–71

Colombo N, Salamon N, Raybaud C, et al (2009) Imaging of malformations of cortical development. Epileptic Disorders: International Epilepsy Journal with Videotape 11(3):194–205. doi:10.1684/epd.2009.0262

Concha L (2014) A macroscopic view of microstructure: Using diffusion-weighted images to infer damage, repair, and plasticity of white matter. Neuroscience 276:14–28. doi:10.1016/j.neuroscience.2013.09.004

Cordero-Grande L, Christiaens D, Hutter J, et al (2019) Complex diffusion-weighted image estimation via matrix recovery under general noise models. NeuroImage 200:391–404. doi:10.1016/j.neuroimage.2019.06.039

Cox RW, Chen G, Glen DR, et al (2017) fmri clustering and false-positive rates. Proceedings of the National Academy of Sciences 114(17):E3370–e3371. doi:10.1073/pnas.1614961114

Ding Z, Morris S, Hu S, et al (2025) Automated whole-brain focal cortical dysplasia detection using mr fingerprinting with deep learning. Neurology 104(11):e213691. doi:10.1212/wnl.0000000000213691

Dong T, Lee HH, Zang H, et al (2025) In vivo cortical microstructure mapping using high-gradient diffusion mri accounting for intercompartmental water exchange effects. NeuroImage 314:121258. doi:10.1016/j.neuroimage.2025.121258

Feizollah S, Tardif CL (2025) 3d mermaid: 3d multi-shot enhanced recovery motion artifact insensitive diffusion for submillimeter, multi-shell, and snr-efficient diffusion imaging. Magnetic Resonance in Medicine 93(6):2311–2330. doi:10.1002/mrm.30436

Fischl B, Dale AM (2000) Measuring the thickness of the human cerebral cortex from magnetic resonance images. Proc Natl Acad Sci USA 97(20):11050 5

Garcia-Hernandez R, Cerdán Cerdá A, Trouve Carpena A, et al (2022) Mapping microglia and astrocyte activation in vivo using diffusion mri. Science Advances 8(21):eabq2923. doi:10.1126/sciadv.abq2923

Genc S, Ball G, Chamberland M, et al (2025) Mri signatures of cortical microstructure in human development align with oligodendrocyte cell-type expression. Nature Communications 16(1):3317. doi:10.1038/s41467-025-58604-w

Gennari AG, Cserpan D, Stefanos-Yakoub I, et al (2023) Diffusion tensor imaging discriminates focal cortical dysplasia from normal brain parenchyma and differentiates between focal cortical dysplasia types. Insights into Imaging 14(1):36. doi:10.1186/s13244-023-01368-y

Ghaderi S, Mohammadi S, Fatehi F (2025) A systematic review of diffusion microstructure imaging (dmi): Current and future applications in neurology research. Brain Disorders 19:100238. doi:10.1016/j.dscb.2025.100238

Gill RS, Lee HM, Caldairou B, et al (2021) Multicenter validation of a deep learning detection algorithm for focal cortical dysplasia. Neurology 97(16):e1571–e1582. doi:10.1212/wnl.0000000000012698

Glenn GR, Helpern JA, Tabesh A, et al (2015) Quantitative assessment of diffusional kurtosis anisotropy. NMR in biomedicine 28(4):448–459. doi:10.1002/nbm.3271

Gross DW, Bastos A, Beaulieu C (2005) Diffusion tensor imaging abnormalities in focal cortical dysplasia. Canadian Journal of Neurological Sciences 32(4):477–482. doi:10.1017/s0317167100004479

Guerrini R, Barba C (2021) Focal cortical dysplasia: an update on diagnosis and treatment. Expert Review of Neurotherapeutics 21(11):1213–1224. doi:10.1080/14737175.2021.1915135

Hansen B (2019) An introduction to kurtosis fractional anisotropy. American Journal of Neuroradiology 40(10):1638–1641. doi:10.3174/ajnr.A6235

Henriques RN, Jespersen SN, Shemesh N (2019) Microscopic anisotropy misestimation in spherical-mean single diffusion encoding mri. Magnetic Resonance in Medicine 81(5):3245–3261. doi:10.1002/mrm.27606

Herberthson M, Boito D, Haije TD, et al (2021) Q-space trajectory imaging with positivity constraints (qti+). NeuroImage 238:118198. doi:10.1016/j.neuroimage.2021.118198

Hui ES, Russell Glenn G, Helpern JA, et al (2015) Kurtosis analysis of neural diffusion organization. NeuroImage 106:391–403. doi:10.1016/j.neuroimage.2014.11.015

Ianuş A, Carvalho J, Fernandes FF, et al (2022) Soma and neurite density mri (sandi) of the in-vivo mouse brain and comparison with the allen brain atlas. NeuroImage 254:119135. doi:10.1016/j.neuroimage.2022.119135

Inverardi F, Chikhladze M, Donzelli A, et al (2013) Cytoarchitectural, behavioural and neurophysiological dysfunctions in the bcnu-treated rat model of cortical dysplasia. The European Journal of Neuroscience 37(1):150–162. doi:10.1111/ejn.12032

Kakita A, Kameyama S, Hayashi S, et al (2005) Pathologic features of dysplasia and accompanying alterations observed in surgical specimens from patients with intractable epilepsy. Journal of Child Neurology 20(4):341–350. doi:10.1177/08830738050200041301

Kawai Y, Aoki I, Umeda M, et al (2010) In vivo visualization of reactive gliosis using manganese-enhanced magnetic resonance imaging. NeuroImage 49(4):3122–3131. doi:10.1016/j.neuroimage.2009.11.005

Kielbinski M, Gzielo K, Soltys Z (2016) Review: Roles for astrocytes in epilepsy: insights from malformations of cortical development. Neuropathology and Applied Neurobiology 42(7):593–606. doi:10.1111/nan.12331

Kleven H, Bjerke IE, Clascá F, et al (2023) Waxholm space atlas of the rat brain: a 3d atlas supporting data analysis and integration. Nature Methods 20(11):1822–1829. doi:10.1038/s41592-023-02034-3

Lampinen B, Zampeli A, Björkman-Burtscher IM, et al (2020) Tensor-valued diffusion mri differentiates cortex and white matter in malformations of cortical development associated with epilepsy. Epilepsia 61(8):1701–1713. doi:10.1111/epi.16605

Lau LA, Dulla CG (2017) Chapter 56 - dysplasias: Cortical freeze lesion. In: Pitkänen A, Buckmaster PS, Galanopoulou AS, et al (eds) Models of Seizures and Epilepsy, 2nd edn. Academic Press, p 845–859, 10.1016/B978-0-12-804066-9.00057-2

Lee C, Kim Y, Kaang BK (2022) The primary motor cortex: The hub of motor learning in rodents. Neuroscience 485:163–170. doi:10.1016/j.neuroscience.2022.01.009

Lee H, Lee HH, Ma Y, et al (2024) Age-related alterations in human cortical microstructure across the lifespan: Insights from high-gradient diffusion mri. Aging Cell 23(11):e14267. doi:10.1111/acel.14267

Lee HH, Papaioannou A, Novikov DS, et al (2020) In vivo observation and biophysical interpretation of time-dependent diffusion in human cortical gray matter. NeuroImage 222:117054. doi:10.1016/j.neuroimage.2020.117054

Lee JW (2016) Frontal focal cortical dysplasias: Too thin here, too thick there, and the folding just isn’t right! Epilepsy Currents 16(4):247–248. doi:10.5698/1535-7511-16.4.247

Lee SK, Kim DI, Mori S, et al (2004) Diffusion tensor MRI visualizes decreased subcortical fiber connectivity in focal cortical dysplasia. NeuroImage 22(4):1826–1829. doi:10.1016/j.neuroimage.2004.04.028

Lerch JP, Pruessner J, Zijdenbos AP, et al (2008) Automated cortical thickness measurements from mri can accurately separate alzheimer’s patients from normal elderly controls. Neurobiology of Aging 29(1):23–30. doi:10.1016/j.neurobiolaging.2006.09.013

Leuze CWU, Anwander A, Bazin PL, et al (2014) Layer-specific intracortical connectivity revealed with diffusion mri. Cerebral cortex 24(2):328–339. doi:10.1093/cercor/bhs311

Li CH, Tam PKS (1998) An iterative algorithm for minimum cross entropy thresholding. Pattern Recognition Letters 19(8):771–776. doi:10.1016/s0167-8655(98)00057-9

Li S, Zheng Y, Sun W, et al (2021) Glioma grading, molecular feature classification, and microstructural characterization using mr diffusional variance decomposition (divide) imaging. European Radiology 31(11):8197–8207. doi:10.1007/s00330-021-07959-x

Lorio S, Adler S, Gunny R, et al (2020) Mri profiling of focal cortical dysplasia using multi-compartment diffusion models. Epilepsia 61(3):433–444. 10.1111/epi.16451

Lundell H, Nilsson M, Dyrby TB, et al (2019) Multidimensional diffusion mri with spectrally modulated gradients reveals unprecedented microstructural detail. Scientific Reports 9(1):9026. doi:10.1038/s41598-019-45235-7

Madan N, Grant PE (2009) New directions in clinical imaging of cortical dysplasias. Epilepsia 50:9–18. doi:10.1111/j.1528-1167.2009.02292.x

Magdoom KN, Pajevic S, Dario G, et al (2021) A new framework for mr diffusion tensor distribution. Scientific Reports 11(1):2766. doi:10.1038/s41598-021-81264-x

McNab JA, Polimeni JR, Wang R, et al (2013) Surface based analysis of diffusion orientation for identifying architectonic domains in the in vivo human cortex. NeuroImage 69:87–100. doi:10.1016/j.neuroimage.2012.11.065

Moroni RF, Inverardi F, Regondi MC, et al (2008) Altered spatial distribution of pv-cortical cells and dysmorphic neurons in the somatosensory cortex of bcnu-treated rat model of cortical dysplasia. Epilepsia 49(5):872–887. doi:10.1111/j.1528-1167.2007.01440.x

Najm I, Lal D, Alonso Vanegas M, et al (2022) The ilae consensus classification of focal cortical dysplasia: An update proposed by an ad hoc task force of the ilae diagnostic methods commission. Epilepsia 63(8):1899–1919. doi:10.1111/epi.17301

Nasser NS, Rajan S, Venugopal Vk, et al (2022) A review on investigation of the basic contrast mechanism underlying multidimensional diffusion mri in assessment of neurological disorders. Journal of Clinical Neuroscience 102:26–35. doi:10.1016/j.jocn.2022.05.027

Nilsson M, Szczepankiewicz F, Brabec J, et al (2020) Tensor-valued diffusion mri in under 3 minutes: an initial survey of microscopic anisotropy and tissue heterogeneity in intracranial tumors. Magnetic Resonance in Medicine 83(2). doi:10.1002/mrm.27959, URL https://onlinelibrary.wiley.com/doi/abs/10.1002/mrm.27959

Novikov DS, Jespersen SN, Kiselev VG, et al (2016) Quantifying brain microstructure with diffusion mri: Theory and parameter estimation. arXiv:161202059 [physics] URL http://arxiv.org/abs/1612.02059

Olesen JL, Østergaard L, Shemesh N, et al (2022) Diffusion time dependence, power-law scaling, and exchange in gray matter. NeuroImage 251:118976. doi:10.1016/j.neuroimage.2022.118976

Palombo M, Ianus A, Guerreri M, et al (2020) Sandi: A compartment-based model for non-invasive apparent soma and neurite imaging by diffusion mri. NeuroImage 215:116835. doi:10.1016/j.neuroimage.2020.116835

Püspöki Z, Storath M, Sage D, et al (2016) Transforms and operators for directional bioimage analysis: A survey. Advances in Anatomy, Embryology, and Cell Biology 219:69–93. doi:10.1007/978-3-319-28549-8_3

Reveley C, Ye FQ, Mars RB, et al (2022) Diffusion mri anisotropy in the cerebral cortex is determined by unmyelinated tissue features. Nature Communications 13(1):6702. doi:10.1038/s41467-022-34328-z

Reveley C, Ye FQ, Leopold DA (2024) Diffusion kurtosis imaging, map-mri and noddi selectively track gray matter myelin density in the primate cerebral cortex. Imaging Neuroscience 2:imag–2–00368. doi:10.1162/imag_a_00368

Robel S, Buckingham SC, Boni JL, et al (2015) Reactive astrogliosis causes the development of spontaneous seizures. The Journal of Neuroscience: The Official Journal of the Society for Neuroscience 35(8):3330–3345. doi:10.1523/jneurosci.1574-14.2015

Rodríguez-Arzate CA, Martínez-Mendoza ML, Rocha-Mendoza I, et al (2021) Morphological and calcium signaling alterations of neuroglial cells in cerebellar cortical dysplasia induced by carmustine. Cells 10(7):1581. doi:10.3390/cells10071581

Rostampour M, Hashemi H, Najibi SM, et al (2018) Detection of structural abnormalities of cortical and subcortical gray matter in patients with mri-negative refractory epilepsy using neurite orientation dispersion and density imaging. Physica Medica 48:47–54. doi:10.1016/j.ejmp.2018.03.005

Schindelin J, Arganda-Carreras I, Frise E, et al (2012) Fiji: an open-source platform for biological-image analysis. Nature Methods 9(7):676–682. doi:10.1038/nmeth.2019

Shemesh N (2018) Axon diameters and myelin content modulate microscopic fractional anisotropy at short diffusion times in fixed rat spinal cord. Frontiers in Physics 6. doi:10.3389/fphy.2018.00049, URL https://www.frontiersin.org/journals/physics/articles/10.3389/fphy.2018.00049/full

Sjölund J, Szczepankiewicz F, Nilsson M, et al (2015) Constrained optimization of gradient waveforms for generalized diffusion encoding. Journal of Magnetic Resonance 261:157–168. doi:10.1016/j.jmr.2015.10.012

Smith SM, Jenkinson M, Woolrich MW, et al (2004) Advances in functional and structural mr image analysis and implementation as fsl. NeuroImage 23 Suppl 1:S208–219. doi:10.1016/j.neuroimage.2004.07.051

Spitzer H, Ripart M, Whitaker K, et al (2022) Interpretable surface-based detection of focal cortical dysplasias: a multi-centre epilepsy lesion detection study. Brain p awac224. doi:10.1093/brain/awac224

Stejskal EO, Tanner JE (1965) Spin diffusion measurements: spin echoes in the presence of a time-dependent field gradient. Journal of Chemical Physics 42(288-292). doi:10.1016/j.jmr.2021.107112

Su TY, Choi JY, Hu S, et al (2024) Multiparametric characterization of focal cortical dysplasia using 3d mr fingerprinting. Annals of Neurology 96(5):944–957. doi:10.1002/ana.27049

Szczepankiewicz F, Hoge S, Westin CF (2019) Linear, planar and spherical tensor-valued diffusion mri data by free waveform encoding in healthy brain, water, oil and liquid crystals. Data in Brief 25:104208. doi:10.1016/j.dib.2019.104208

Szczepankiewicz F, Westin CF, Nilsson M (2021) Gradient waveform design for tensor-valued encoding in diffusion mri. Journal of Neuroscience Methods 348:109007. doi:10.1016/j.jneumeth.2020.109007

Topgaard D (2019) Diffusion tensor distribution imaging. NMR in biomedicine 32(5):e4066. doi:10.1002/nbm.4066

Tournier JD, Smith R, Raffelt D, et al (2019) Mrtrix3: A fast, flexible and open software framework for medical image processing and visualisation. NeuroImage 202:116137. doi:10.1016/j.neuroimage.2019.116137

Urquia-Osorio H, Pimentel-Silva LR, Rezende TJR, et al (2022) Superficial and deep white matter diffusion abnormalities in focal epilepsies. Epilepsia 63(9):2312–2324. doi:10.1111/epi.17333

Villaseñor PJ, Cortés-Servín D, Pérez-Moriel A, et al (2023) Multitensor diffusion abnormalities of gray matter in an animal model of cortical dysplasia. Frontiers in Neurology 14. URL https://www.frontiersin.org/articles/10.3389/fneur.2023.1124282

Vis G, Nilsson M, Westin CF, et al (2021) Accuracy and precision in super-resolution mri: Enabling spherical tensor diffusion encoding at ultra-high b-values and high resolution. NeuroImage 245:118673. doi:10.1016/j.neuroimage.2021.118673

Walger L, Schmitz MH, Bauer T, et al (2025) A public benchmark for human performance in the detection of focal cortical dysplasia. Epilepsia Open 10(3):778–786. doi:10.1002/epi4.70028

Westin CF, Knutsson H, Pasternak O, et al (2016) Q-space trajectory imaging for multidimensional diffusion mri of the human brain. NeuroImage 135:345–362. doi:10.1016/j.neuroimage.2016.02.039

Winston GP, Micallef C, Symms MR, et al (2014) Advanced diffusion imaging sequences could aid assessing patients with focal cortical dysplasia and epilepsy. Epilepsy Research 108(2):336–339. doi:10.1016/j.eplepsyres.2013.11.004

Wong M (2009) Animal models of focal cortical dysplasia and tuberous sclerosis complex: Recent progress toward clinical applications. Epilepsia 50:34–44. doi:10.1111/j.1528-1167.2009.02295.x

Zhang H, Schneider T, Wheeler-Kingshott CA, et al (2012) Noddi: Practical in vivo neurite orientation dispersion and density imaging of the human brain. NeuroImage 61(4):1000–1016. doi:10.1016/j.neuroimage.2012.03.072

Zhu A, Michael ES, Li H, et al (2025) Engineering clinical translation of ogse diffusion mri. Magnetic Resonance in Medicine 94(3):913–936. doi:10.1002/mrm.30510

